# ER-Mitochondria Contact Sites expand during mitosis

**DOI:** 10.1101/2023.06.22.546089

**Authors:** Fang Yu, Raphael Courjaret, Asha Elmi, Ayat Hammad, Melanie Fisher, Mark Terasaki, Khaled Machaca

## Abstract

Membrane contact sites between various organelles define specialized spatially defined signaling hubs, they are of great interest to better understand inter-organelle communication and its implications on cellular physiology. ER-mitochondria contact sites (ERMCS) are one of the best studied and mediate Ca^2+^ signaling that regulates mitochondrial bioenergetics. However, little is known about ERMCS during mitosis. Here we show that ERMCS expand during mitosis using transmission electron microscopy, serial EM coupled to 3D reconstruction, and ERMCS markers. ERMCS expansion in mitosis is functionally significant as it is associated with enhanced Ca^2+^ coupling between the ER and mitochondria resulting in heightened activation of mitochondrial dehydrogenases. Our data suggest that ERMCS remodeling in mitosis is important to meet the increased energy needs during cell division.

## Introduction

Contact sites are close appositions between cellular organelles that are emerging as specialized signaling hubs for non-vesicular lipid transfer and Ca^2+^ signaling among other functions (Scorrano et al., 2019). One of the earliest investigated organelle contact sites is between the ER and mitochondria (ERMCS) and has been shown to facilitate redox and Ca^2+^ signaling between the two organelles to regulate cellular metabolism (Csordas et al., 2018). Although ERMCS have been heavily investigated their regulation during cell division remains obscure.

The process of cell division must ensure equal distribution of intracellular organelles to support the survival and functionality of daughter cells. This could be a passive process if the organelle is evenly distributed throughout the cytoplasm, but for most organelles including the mitochondria this is not the case. So typically, mitochondria fragment during the G2/M transition of the cell cycle to support equal partitioning to the daughter cells (Carlton et al., 2020; Labbe et al., 2014). Mitochondria present an additional complexity compared to other organelles as they contain their own DNA that needs to replicate in parallel with the nuclear DNA before cell division. This is coupled to mitochondrial fusion and elongation during G1/S and fragmentation during G2/S (Mishra and Chan, 2014). Interestingly, both mitochondrial fission and fusion are initiated at ERMCS (Bleazard et al., 1999; Ferguson and De Camilli, 2012; Korobova et al., 2013). In addition, mitochondria are the energy source for the cell, so they must maintain their functionality as the energy demands during cell division are elevated (Salazar-Roa and Malumbres, 2017; Wang et al., 2014; Zhao et al., 2019).

Mitochondrial fission is regulated by the dynamin-related GTPase DRP1, which localizes to ER-mitochondria contact sites (Labbe et al., 2014). The cell cycle kinases Aurora A and CDK1 phosphorylate DRP1 in mitosis resulting in increased mitochondrial fission and thus fragmentation in preparation for cytokinesis (Kashatus et al., 2011). Mitochondrial respiration is increased during mitosis through CDK1 phosphorylation of different subunits in complex I (CI) (Wang et al., 2014). In addition, a Ca^2+^ transient in metaphase has been shown to be important for maintaining cellular energy levels in mitosis through an MCU-dependent process, showing that mitochondrial Ca^2+^ uptake is important for mitosis progression (Zhao et al., 2019).

Similar to mitochondria the ER as well remodels during mitosis (Lu et al., 2009), however whether ERMCS connectivity is affected in mitosis compared to interphase remains unclear. Here we show using multiple approaches that ERMCS expand during mitosis and that this expansion is functionally significant as it is associated with more efficient ER-mitochondrial Ca^2+^ coupling resulting in enhanced activation of matrix dehydrogenases.

## Methods

### Cell culture

HEK293, Hela, and Jurkat cells were obtained from ATCC. HEK293 and Hela cells were cultured in DMEM media (Invitrogen) supplemented with 10% FBS (Sigma), 100U/ml penicillin, and 100μg/ml streptomycin. Jurkat cells were cultured in RPMI 1640 media (Invitrogen) supplemented with 10% FBS (Sigma), 100U/ml penicillin and 100μg/ml streptomycin. Cells were incubated at 37°C and under 5% CO_2_ for the indicated time. For transmission electron microscopy (TEM), Ca^2+^ imaging and NAD(P)H imaging Jurkat cells were untreated or treated with 100 ng/ml of Nocodazole (Sigma) for 18 h for mitotic synchronization.

The stable HEK293 and Hela cell lines with ER-mito MCS reporter expression were generated by lentiviral infection. Briefly, cells were infected with pLVX -Mitot-spGFP11×2 and plx304-spGFP1-10-ERt using standard procedure described elsewhere (Yang et al., 2018). Infected cells were then selected with puromycin and blasticidin and cells with moderate expression levels were enriched by a FACSAria II cell sorter (BD Bioscience).

### Flow cytometry

Mitochondrial volume was measured by loading Jurkat cells with 100 nM Mitotracker red CMXRox (Invitrogen) according to the manufacturer’s instructions. Cells were harvested with Fixable Viability Dyes eFluor 506 (eBioscience) prior to fixation. Fixed and permeabilized cells were subjected to immunostaining with Alexa Fluor 647 Rat anti-Histone H3 (pS28) (Clone HTA28) antibody, a well-known marker for mitosis, according to the manufacture instruction (BD Biosciences). The measurement of mitochondrial volume was conducted using a BD Fortessa X20 cell analyzer. Viable naturally occurring mitotic and interphase cells were analyzed. Flow cytometry data were analyzed by the FlowJo software (BD).

### Confocal Microscopy

Cells were cultured on 35-mm glass-bottom dishes (MatTek). Cells were stained at 37°C for 20 minutes in culture medium with 100 nM Mitotracker deep red to indicate mitochondria and stained either Hoechst33324 (Invitrogen) or NuclearID red (Enzo Life Sciences) to distinguish interphase and mitosis. To Quantify ERMCS in interphase and mitotic cells, Cells were then fixed with 4% PFA in PBS for 10 minutes at room temperature. Interphase and naturally occurring mitotic cells were imaged using a Zeiss LSM 880 confocal microscope with Airyscan using a Plan Apo 63×/1.4 oil DIC II objective with pinhole 1AU. Images were analyzed using ZEN Black software.

### TEM and 3D reconstruction

To prepare samples for TEM, Jurkat cell pellets were treated with a modified Karmovsky’s fix and followed by a secondary fixation in reduced osmium tetroxide as described previously (Yu et al., 2019). The samples were then En bloc stained with uranyl acetate and dehydrated with graded ethanol before being embedded in an Epon analog resin. The 65 nm ultrathin sections were contrasted with lead citrate and imaged using a JEM 1400 electron microscope (JEOL, USA, Inc.) operating at 100 kV. Digital images were captured using a Veleta 2K × 2K CCD camera (Olympus-SIS). The length, density, gap of ERMCS, and perimeter of mitochondria in each cell were measured and calculated by ImageJ. The density of ERMCS was quantified as ERMCS length divided by the circumference of the mitochondrion.

For 3D reconstruction from serial sections, the methods were the same as used previously (Terasaki et al., 2013). Briefly, 50 nm thick sections were collected on kapton tape using an ATUM (automated tape collecting microtome), then imaged by back scatter in a scanning electron microscope. Images were aligned in FIJI/Image J and ER contact sites were traced in Reconstruct (version 1.1).

### Ca^2+^ and NAD(P)H imaging

Imaging of Jurkat cells was performed using a PTI Easy Ratio Pro system (software version 1.6.1.0.101; Horiba Scientific) composed of a DeltaRAMX monochromator and a CoolSnapHQ^2^ camera attached to an Olympus IX71 inverted microscope fitted with a 20x/0.75 lens. For intracellular Ca^2+^ imaging the cells were loaded for 30 min with 2 mM Fura2-AM in a Ca^2+^- containing media (composition below) at room temperature and washed once in Ca^2+^-containing or Ca^2+^-free media prior to the experiments. Excitation was performed at 340 nm and 380 nm for a duration of 100 ms at a 0.1 Hz frame rate and the ratio of the fluorescence intensity at 340/380 measured. For NAD(P)H measurement the cells were excited at 340 nm for 500 ms at 0.1 Hz and emitted light collected through a 420 nm long pass filter (Olympus UMWU2). Jurkat cells were plated on Poly-D-lysine coated coverslips (MatTek) and left for 10 min to allow adhesion to the coverslip. The cells were activated using 1.0 μg/ml of BioLegend Ultra-LEAF™ Purified anti-human CD3 Antibody (clone OKT3) and anti-human CD28 Antibody (clone 28.2). The extracellular media contained (in mM): NaCl 1.5, KCl 5, CaCl_2_ 1.5, D-glucose 10, MgCl_2_ 1.5 HEPES 20, pH 7.4. Image analysis was performed using Image J.

### Statistical Analyses

Data are presented as mean ± SEM. Groups were compared using the Prism 9 software (GraphPad) using the statistical tests indicated in the figure legend. Statistical significance is indicated by p values in the Figure legends.

## Results and Discussion

### ERMCS expand in mitosis

To assess ERMCS during mitosis at the ultrastructural level, we chose Jurkat T lymphocytes due to their relatively less complex intracellular organelle organization. ERMCS are readily detected in both interphase and mitotic cells in thin section electron micrographs (Fig. 1A and 1B). Mitosis is associated with dramatic remodeling of organelles: nuclear envelope breakdown, chromosome condensation, and ER remodeling (Fig. 1B). The ER typically forms a ring at the cell cortex in mitotic Jurkat T-cells (Fig. 1B), and we have previously shown that ER-PM contact sites (ERPMCS) are downregulated significantly during mitosis (Yu et al., 2019). ERMCS also appear to remodel during mitosis but in the opposite direction, where there is an increase in the length of the contact between the ER and mitochondria (Fig. 1B). In some cases, the ER is in close apposition through the entire length of a mitochondrion (Fig. 1B, lower right panel). Furthermore, in thin sections mitochondria appear to have a smaller diameter in mitosis, consistent with their previously documented fragmentation (Fig. 1B)

**Figure 1.**
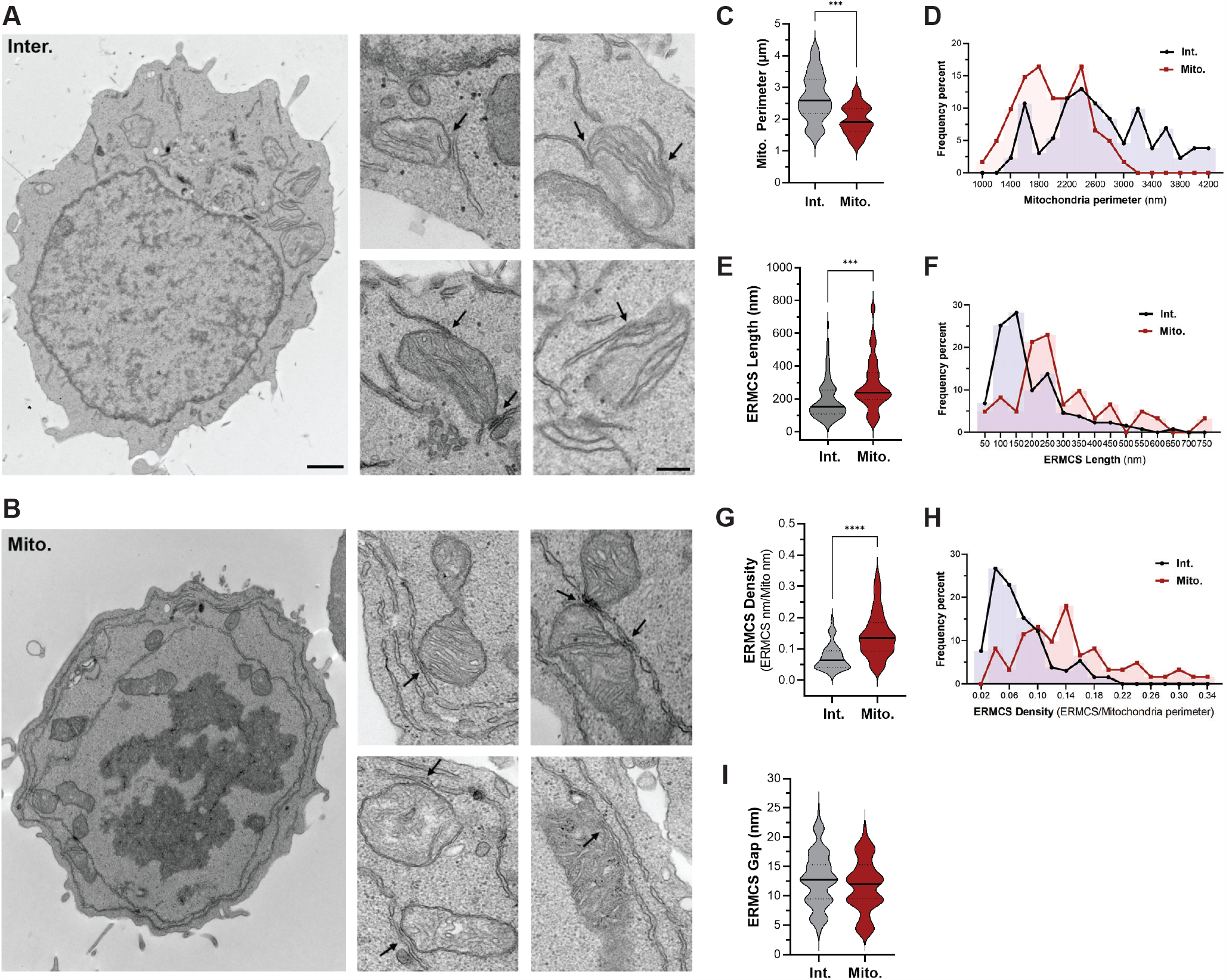
TEM analysis of ERMCS in interphase (Inter) and mitotic (Mito) Jurkat cells. (**A-B**). Representative TEM images from Inter (**A**) and Mito (**B**) cells. Black arrows indicate the ERMCS defined as close contacts between the ER and mitochondrion at <30 nm. A representative whole cell is shown at low magnification (scale bar 1 μm), with several zoomed-in regions (scale is 250 nm) containing ERMCS (arrows in A and B). (**C, E, G, I**) Quantification of mitochondrial perimeter (**C**), ERMCS length (**E**), ERMCS density (**G**), and ERMCS gap (**I**) in interphase and mitotic cells (Mean ± SEM, unpaired two tailed t-test, n=61-131, *** p≤0.001. **** p≤0.0001). (**D, F, H**) Frequency histogram of mitochondrial perimeter (**D**), ERMCS length (**F**), and ERMCS density (**H**).

Quantification of mitochondrial perimeter from EM images sections confirms their fragmentation during mitosis with a smaller average perimeter (Fig. 1C) and a shift in the histogram distribution of the population to the left (Fig. 1D). Furthermore, ERMCS length increases significantly during mitosis (Fig. 1E) with a shift in the histogram toward ERMCS above 200 nm in length as compared to interphase ERMCS whose distribution shows a peak in the 100-150 nm range (Fig. 1F). The increase in length is supported by increased ERMCS mitochondrial density in mitosis (Fig. 1G and 1H). We defined ERMCS density as the percent of the mitochondrial perimeter occupied by ERMCS. Increased ERMCS length coupled mitochondrial fragmentation explain the observed enhanced ERMCS density (Fig. 1G and 1H). There was no difference in the ERMCS gap distance in mitosis as compared to interphase (Fig. 1I). Furthermore, although mitochondria fragment during mitosis, the total mitochondrial volume, assessed using Mitotracker staining, was not altered during mitosis (Fig. 2A).

**Figure 2.**
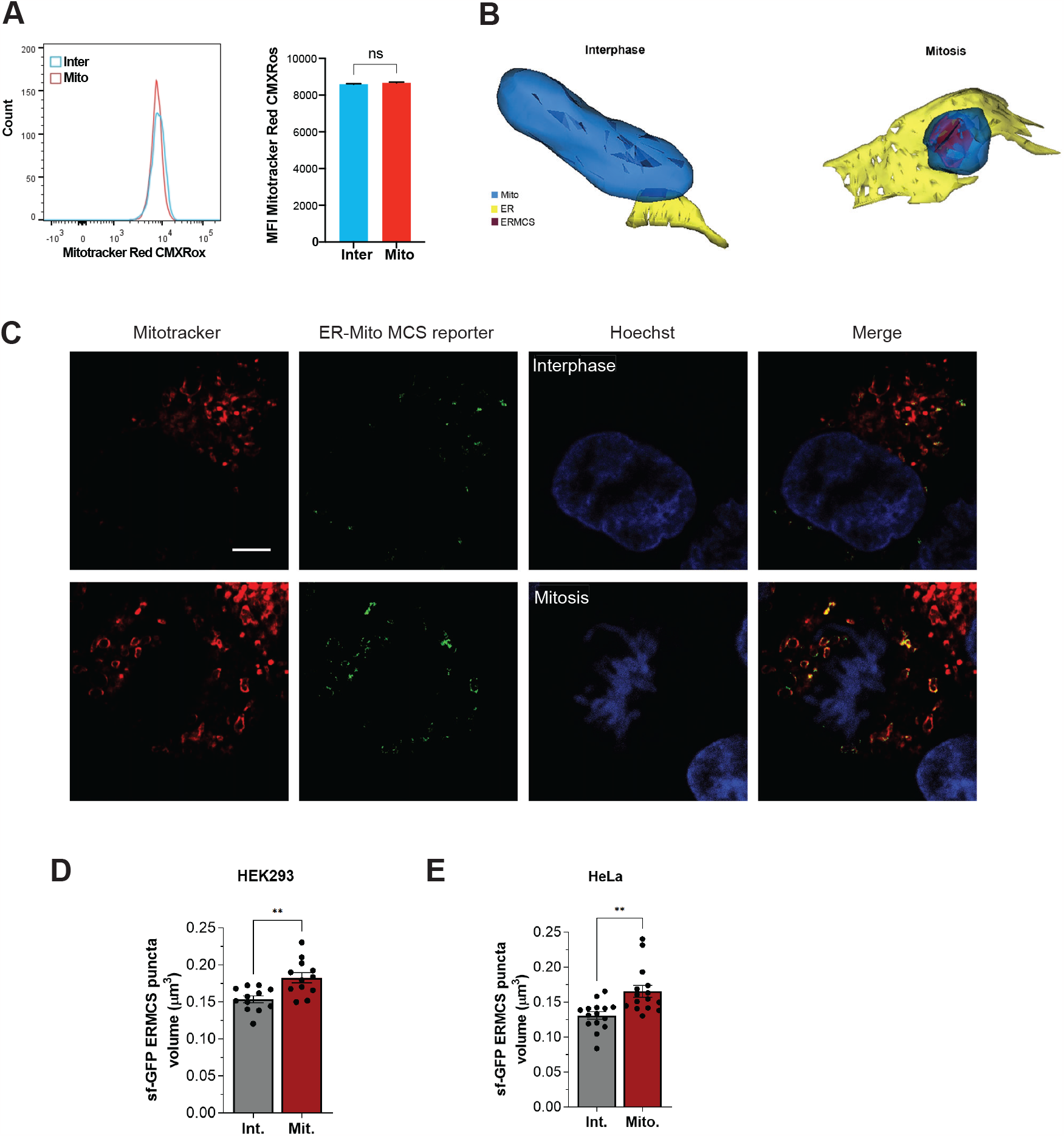
ERMCS increase in mitotic HEK293 and Hela cells. (**A**) Mitochondrial volume in interphase (Inter) and naturally occurring mitotic (Mito) Jurkat cells was measured using MitoTracker Red. Representative flow cytometry plots (left) and summary data (right) are shown. Mean ± SEM of 4 repeat experiments. (**B**) Representative 3D reconstruction in interphase and mitosis from serial EM of ER (yellow) and mitochondria (blue) with ERMCS highlighted in red. (**C**) Representative confocal images of HEK293 cells stable expressing the ERMCS reporter either in interphase or naturally occurring mitosis. Cells were stained with Mitotracker deep red to identify mitochondria and Hoechst to differentiate cells in interphase or mitosis. Scale bar 5 μm. (**D-E**). Quantification of ERMCS volume using Imaris software after 3D reconstruction of confocal stacks in HEK293 (D) and HeLa (E) cells stably expressing the split GFP ERMCS reporter. (**D**) Mean ± SEM, unpaired two tailed t-test, n=12, p=0.0019. (**E**) Mean ± SEM, unpaired two tailed t-test, n=15, p=0.0017.

To expand the characterization of ERMCS remodeling we performed three-dimensional reconstruction of ERMCS in both interphase and mitosis from serial EM sections of Jurkat T cells. We performed the serial EM sectioning on three cells each in interphase and mitosis and performed 3D reconstruction by direct visual inspection as detailed in Methods. Examples of two such reconstructions are shown in Figure 2B and confirm the significant expansion of ERMCS in mitosis, where the ER cups the mitochondrion resulting in a larger ERMCS interphase.

### ERMCS are enhanced in mitosis in different cell types

T cells are ideal to analyze ERMCSs at the ultrastructural level due to their small size, large volume occupied by the nucleus, and relatively low organellar complexity. However, T cells are fairly specialized cells, so we wondered whether the increase in ERMCS in mitosis is a universal process or may be limited to T cells. To address this question, we resorted to a higher throughput approach to assess ERMCS using the split GFP reporter (sGFP) with two reporters each containing a complementary GFP fragment expressed in the ER and mitochondrial membrane, so functional GFP would be formed only if the two membranes are in close enough proximity at ERMCS (Yang et al., 2018). We generated stable cell lines expressing the sGFP-ERMCS reporter at low levels in both HeLa and HEK293 cells, and co-stained with Hoechst to allow identification of mitotic cells and mitotracker to confirm localization to the mitochondria (Fig. 2C). As observed in Jurkats by EM, ERMCS increase in mitosis (Fig. 2C). Quantification of the GFP-positive ERMCS volume showed a significant increase in mitotic cells in both HEK293 (Fig. 2D) and HeLa cells (Fig. 2E).

### ERMCS expansion in mitosis is associated with enhanced Ca^2+^ transfer to mitochondria

As discussed above mitochondrial respiration has been shown to increase in mitosis (Wang et al., 2014; Zhao et al., 2019) presumably to meet the increased energy demands of the dividing cell. The expansion of ERMCS could contribute to supporting increased mitochondrial respiration especially that mitochondrial Ca^2+^ transients have been shown to be important to support the cell’s energy demands in mitosis. Given the close apposition of the two organelles membranes at ERMCS, they are preferred sites for efficient Ca^2+^ transfer between the ER to activate Ca^2+^- sensitive matrix dehydrogenases (Csordas et al., 2010; Katona et al., 2022). Furthermore, the MCU channel in the inner mitochondrial membrane has been shown to be important for the mitochondrial Ca^2+^ transients in mitosis (Zhao et al., 2019).

Our data so far show that ERMCS expand during mitosis, arguing that this expansion could support better Ca^2+^ coupling between the ER and mitochondria. To test whether this is the case, we measured Ca^2+^ signals and NAD(P)H autofluorescence as a readout of mitochondrial dehydrogenase activity (Katona et al., 2022) (Fig. 3). Cross linking the TCR using anti-CD3 and anti-CD28 antibodies in Jurkat T cells stimulated Ca^2+^ signals that were of significantly higher amplitude in interphase compared to mitotic cells arrested in metaphase using nocodazole (Fig. 3A and 3B). We recorded in parallel NAD(P)H autofluorescence in interphase (Fig. 3C) and mitotic cells (Fig. 3D). NAD(P)H autofluorescence was similar following TCR stimulation in interphase and mitotic cells (Fig. 3E), despite the significantly reduced Ca^2+^ transient in mitosis (Fig. 3B).

**Figure 3.**
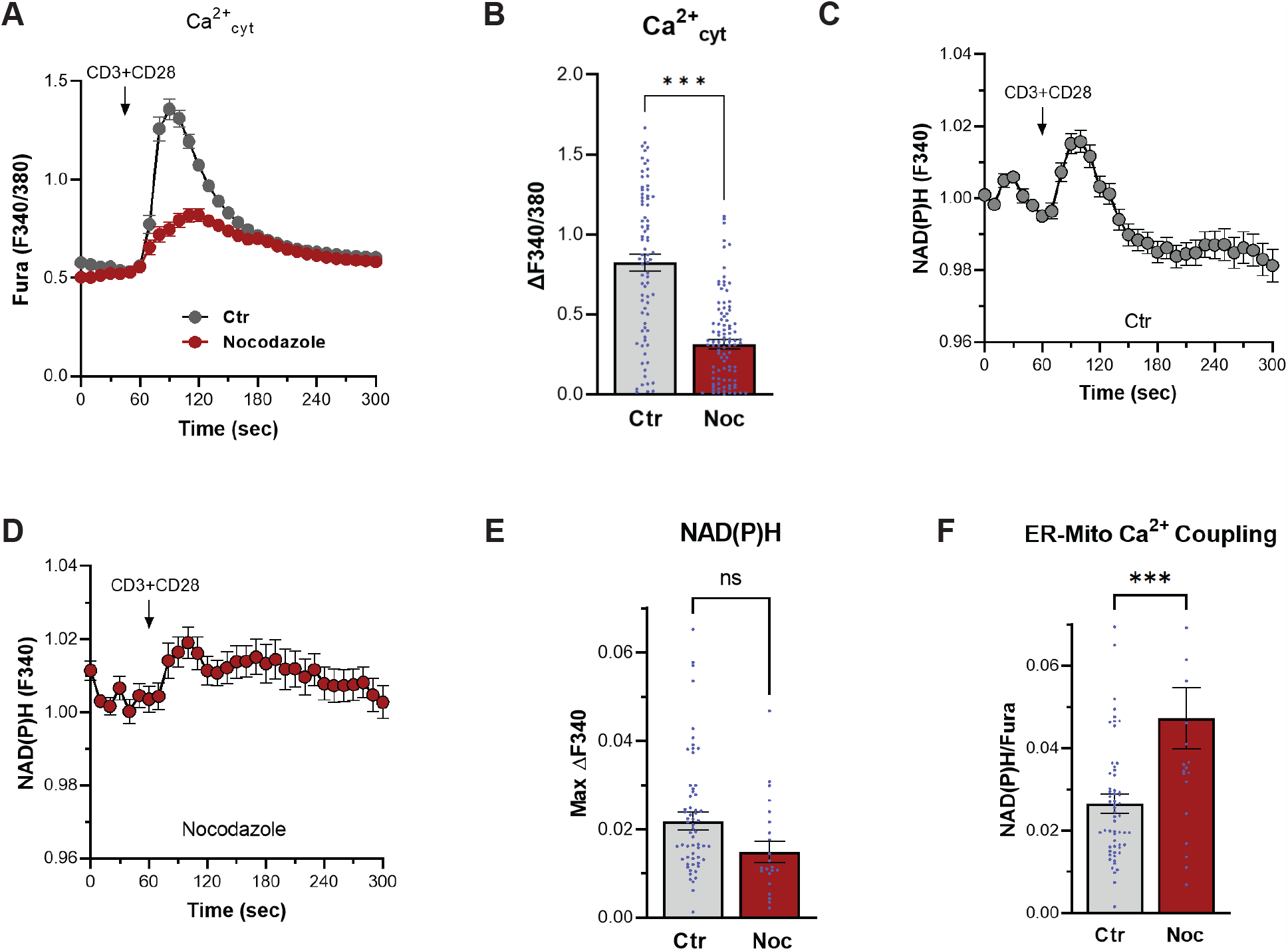
Intracellular Ca^2+^ signals and NAD(P)H levels in interphase and mitosis. **(A)** Cytoplasmic Ca^2+^ (Ca^2+^_cyt_) time course in response to CD3/CD28 stimulation in Jurkat T cells in asynchronous or mitosis (nocodazole). **(B)** Quantification of the peak Ca^2+^_cyt_ response. Mean ± SEM, unpaired two tailed t-test, n=79-104, p<0.001=0.0017. **(C-D)** Time course of the NAD(P)H autofluorescence (measured at 340 nm) induced by CD3/CD28 stimulation is asynchronous (C) and mitotic (D) cells. **(E)** Quantification of the peak NAD(P)H autofluorescence. Mean ± SEM, unpaired two tailed t-test, n=23-61. **(F)** Coupling between the ER and the mitochondria quantified as the ratio between the peak (NAD(P)H signal and the maximum Ca^2+^_cyt_ induced by CD3/CD38 stimulation. Mean ± SEM, unpaired two tailed t-test, n=23-61, p=0.0008.

We assessed ER-Mitochondrial Ca^2+^ coupling by quantifying the ratio of the NAD(P)H autofluorescence signal to that of the Ca^2+^ signal (Fig. 3F). ER-Mito Ca^2+^ coupling was significantly enhanced in mitosis, where matrix dehydrogenases were stimulated more effectively in response to a smaller Ca^2+^ transient (Fig. 3F). This argues that the expansion of ERMCS in mitosis is functionally important to allow mitochondria to more effectively uptake Ca^2+^ to stimulate respiration.

Our results show that ERMCS expand during mitosis and that this expansion is associated with more efficient transfer of Ca^2+^ between the ER and mitochondria. During mitosis, the cell undergoes dramatic changes, including organelle remodeling, chromosome condensation, and nuclear envelope breakdown to name a few. These processes are energy intensive and thus require functioning mitochondria to fuel them during mitosis. The expansion of ERMCS in mitosis is one mechanism that supports mitochondrial respiration in mitosis.

## Additional Information

### Data Availability Statement

All data reported are included in the manuscript proper and are available on request.

## Acknowledgments

We are grateful to Microscopy Core at WCMQ and the EM Core at WCM for their support in multiple experiments.

